# The Role of Lrig1 in the Development of the Colonic Epithelium

**DOI:** 10.1101/2023.05.02.539114

**Authors:** Rachel E. Hopton, Nicholas J. Jahahn, Anne E. Zemper

**Author notes:** Corresponding Author: Anne E. Zemper, Ph.D. 1229 University of Oregon, 1318 Franklin Blvd. Room 273 Onyx Bridge Eugene, OR 97403, ORCID: 0000-0001-8238-1406. **NEW & NOTEWORTHY:** Our studies define the importance of studying Lrig1 and its role in colon development. We address a critical gap in the intestinal development literature and provide new information about the molecular cues that guide colon development. Using a novel inducible knockout of Lrig1, we show Lrig1 is required for appropriate colon epithelial growth and illustrate the importance of Lrig1 in the establishment of developing colonic crypts.

## Abstract

Growth and specification of the mouse intestine occurs in utero and concludes after birth. While numerous studies have examined this developmental process in the small intestine, far less is known about the cellular and molecular cues required for colon development. In this study, we examine the morphological events leading to crypt formation, epithelial cell differentiation, areas of proliferation, and the emergence and expression of a stem and progenitor cell marker Lrig1. Through multicolor lineage tracing, we show Lrig1 expressing cells are present at birth and behave as stem cells to establish clonal crypts within three weeks after birth. In addition, we use an inducible knockout mouse to eliminate Lrig1 during colon development and show loss of Lrig1 restrains proliferation within a critical developmental time window, without impacting colonic epithelial cell differentiation. Our study illustrates the morphological changes that occur during crypt development and the importance of Lrig1 in the developing colon.

## INTRODUCTION

Over the last 50 years our understanding of the origin, specification and patterning of model mammalian organs and organ systems has exponentially increased (1–3). For digestive organs, these data have contributed to our comprehension of human development (2, 4–6), informed our knowledge of diseases etiology (2, 6–8), and in some cases, have also facilitated the development of therapeutic interventions (9–11). These data sets describing the molecular requirements for growth and development of the small intestine; however, mammalian colon development is less well understood. A handful of studies from the last 30-40 years (3, 12–20) provide the groundwork for our current understanding of colonic development, yet the molecular mechanisms that govern colon development from birth to adulthood remain largely unknown.

In the mouse, the majority of the growth and specification of the colon occurs postnatally. In the immediate days and weeks following birth, the colon elongates, and expand to form functional crypt units (12–23). During this timeframe, three functionally distinct regions of the colon are also developing. The region adjacent the cecum is commonly referred to as the “proximal” colon and harbors distinct mucosal folds not present in the “middle” and the “distal” colon, located posterior to the proximal region. The distal colon grows from a flat nascent mucosal layer, forming small invaginations on the surface by the second postnatal day (14). These invaginations form the *bona fide* precursors to adult colonic crypts by P7 (13); however, the molecular characteristics of these nascent crypts have not been defined.

In the small intestine, cell proliferation and specification is directed by a heterogeneous population of intestinal stem cells (13, 24, 25). These stem cells reside at the base of the colonic crypts and divide to create a progenitor cell pool known as transit-amplifying (TA) cells. TA cells further divide, move up the crypt in a conveyor belt-like fashion, and become more differentiated, committing to either an absorptive or secretory colonic epithelial cell fate (Barker et al., 2014). Over recent years, numerous molecular markers of these stem, TA and differentiated cells have been discovered (18, 25–27). Despite these advances, very little has been done to characterize these markers during colon development (23, 28) and even less is known about what functional role, if any, these proteins play in crypt development.

The recent expansion in our understanding of adult colon stem cell behavior and the biology of colonic crypts has largely been facilitated by genetically labeling the cells in the stem cell compartment utilizing several lineage-tracing approaches (18, 25, 29–32). These experimental tactics have provided the basis for our understanding that adult colonic crypts are clonal units, which arise from a single stem cell population at the base of each crypt. Due to their clonal nature, any genetic or epigenetic changes in colon stem cells can persist and are retained in the daughter cell population (15, 33, 34). Studies examining human genetic and epigenetic variations in crypts demonstrate that aberrant crypts can arise from mutant stem cell populations, and have a widespread, deleterious impact along the length of the adult colon (35, 36). Clonality is a defining tenet of colon crypts (25, 34), yet it is unknown when clonality is established during crypt development.

One important molecular feature of adult intestinal and colonic crypts is the expression of a protein called Leucine-rich repeats and immunoglobulin-like domains 1 (Lrig1) (17, 18). Lrig1 is a transmembrane protein which decorates the surface of stem and progenitor cells in the adult colon and functions as a tumor suppressor (18). Lrig1 is required for proper intestinal development; complete deletion of *Lrig1* can result in premature postnatal death of mice, with digestive tract abnormalities (17). In addition, after acute injury to the epithelium, Lrig1 expression is rapidly expanded in the colon as a part of the regenerative process and lineage tracing from Lrig1+ cells demonstrates the cells proliferate and divide to replenish colonic crypts (18, 21, 37). Lrig1 is important for mouse development, as Lrig1-expressing (Lrig1+) cells aid in regulating a balance of proliferation and differentiation in the developing brain and skin (38–40). Lrig1 is also present in the gastrointestinal tract at birth (18), but the role of Lrig1 and Lrig1-expressing cells have not been defined.

The goal of our study was to identify key cellular, molecular, and morphological features of colon development using a loss-of-function approach. We first defined the morphological and molecular characteristics of wildtype, distal colonic crypt development from birth to three-weeks of age. Next, we examined the role of Lrig1+ cells. To achieve this, we lineage traced Lrig1+ cells using a multicolor reporter during colon development. Further, to test the requirement for Lrig1 in colon development, we utilized CreERT2/LoxP (18) (EUCOMM) mice to inducibly eliminate Lrig1 at birth and examined the impact in the first three weeks of life. Our data indicate Lrig1+ cells are present at P1 and give rise to clonal crypts three weeks later. We show Lrig1 is not required for crypt cell differentiation but is required to suppress proliferation during colonic crypt development two weeks after birth.

## METHODS

### Mice

Mice were housed in a specific pathogen-free environment under strictly controlled light cycle conditions, fed a standard rodent lab chow and provided water *ad libitum*. All procedures were approved and performed in accordance with the University of Oregon Institutional Animal Care and Use Committee. *C57BL/6* and *Lrig1^tm1.1(cre/ERT2)Rjc^/J* (*Lrig1-CreERT2*) mice (18) were obtained from Jackson Laboratory (Bar Harbor, ME). Lrig1^tm1a(EUCOMM)Wtsi^ (*Lrig1-flox/flox*) mice (EUCOMM) were generously provided by Robert J. Coffey (Vanderbilt University Medical Center). *Gt(ROSA)26Sor^tm1(CAG-Brainbow2.1)Cle^/J* mice (*R26R-Confetti*) (34) were generously provided by Kryn Stankunas (University of Oregon). *Lrig1-CreERT2/+* mice were crossed to *R26R-Confetti* mice or *Lrig1-flox/floxl* mice. Mice were sacrificed between P1 and 6 weeks of age. For multicolor lineage labeling, *Lrig1-CreERT2/+* mice were crossed to *R26R-Confetti* mice; resulting *Lrig1-CreERT2/+;R26R-Confetti* mice were injected via intraperitoneal (IP) injection once with 30mg/kg tamoxifen (Sigma-Aldrich, T5648, suspended in corn oil) at P1 and the colon was harvested at P5, P17, and P22. For Cre-mediated deletion of Lrig1, *Lrig1-CreERT2/+* mice were crossed to *Lrig1-flox/flox* mice and the resulting *Lrig1-CreERT2/ flox* mice and *Lrig1-WT/ flox* (littermate controls) were injected once with 30mg/kg tamoxifen at P5 and the colon was harvested at P7, P14, P21 and at 6 weeks (P42).

### Tissue preparation for staining

Colons were dissected from the mice and flushed with ice-cold Phosphate Buffered Saline (PBS, Grow Cells, MRGF-6395) and ice-cold 4% Paraformaldehyde (PFA; Sigma, 158127)/PBS then pinned as a tube or placed into 6-well tissue culture dishes (Falcon, 353046). Colon was then fixed in 4% PFA/PBS for 20 minutes (P1-P7) or 60 minutes (P13-P42) at room temperature with light oscillation. For frozen sections, fixed colons were washed 3×5 minutes in PBS and incubated in 30% sucrose overnight at 4°C, then blocked in Optimal Cutting Temperature Compound (OCT, Tissue-Tek® Sakura, 4583) as tubes, separating the proximal, middle, and distal portions of the colon. Frozen blocks were placed at −80°C for storage. For paraffin sections, fixed colons were washed 3×5 minutes in PBS, incubated in 70% ethanol overnight at room temperature followed by serial ethanol dehydrations and subsequent xylene clearing washes prior to paraffin embedding. Paraffin blocks were stored at room temperature.

### Immunofluorescence and Histology

For both frozen and paraffin preparations, 10μm transverse sections of colon were placed on Superfrost™ Plus (Fisher Scientific, 12-550-15) or Millennia Command™ slides (StatLab, MCOMM). For frozen sections, slides were thawed 15 minutes at room temperature, washed 3×3 minutes in PBS, blocked in blocking buffer composed of 1% Bovine Serum Albumin (BSA), 1M CaCl_2_, and 0.03% Triton X-100 (Sigma, T8787) for 1 hour before primary antibody was added. All primary antibodies were suspended in blocking buffer. All stains were performed using an overnight incubation at 4°C, except for Ki-67 staining in Figure 5D, D’, D’’’, which was incubated overnight at room temperature. The following primary antibodies were used for frozen sections: anti-Lrig1 (R&D, AF3688 1:250), anti-Non-phospho (Active) β-Catenin (Ser45) (D2U8Y) XP (Cell Signaling Technology, 19807 1:1000), anti-Ki-67 (Invitrogen, 14-5698-82, 1:100), anti-Chromogranin A (LK2H10) (AbD Serotec, MCA845 1:200), anti-DCAMKL1 (Abcam, ab31704, 1:1000), anti-Alexa Fluor™ 488 Phalloidin 488 (Invitrogen, A12379, 1:2000). On the following day, slides were washed 3×3 minutes in PBS then stained with secondary antibody (Jackson ImmunoResearch, 1:500) diluted in the blocking buffer for 1 hour at room temperature. For double staining of Ki-67 and Lrig1, the Ki-67 antibody stain was first accomplished, followed by a one-hour incubation with Lrig1 antibody at room temperature. The secondary antibody staining protocol described above. For phalloidin staining, slides were washed and blocked as described above and the conjugated antibody was incubated for 1 hour at room temperature. Prior to coverslipping, slides were washed 3×3 minutes in PBS, counterstained with DAPI, and mounted with N-propyl gallate/glycerol mounting medium. For Muc2 (anti-Muc2 Cloud-Clone, PAA705Mu01, 1:100), paraffin-embedded tissue sections underwent conventional deparaffinization in xylenes and rehydration via ethanol washes and water. Antigen retrieval was performed using a 1x Citrate Buffer (Sigma, C9999) in a pressure cooker for 15 minutes on high, followed by quick release and slides placed on ice. Slides were blocked in blocking buffer described above with the addition of 10% donkey serum for 1 hour before addition of the primary antibody. Primary antibody was diluted in PBS and slides were incubated overnight at 4°. On the following day, slides were washed 3×3 minutes in PBS then stained with secondary antibody (Jackson ImmunoResearch, 1:500) diluted in PBS for 1 hour at room temperature. To assess colon morphology, paraffin-embedded tissue sections were stained with Hematoxylin and Eosin (VWR, 95057) using standard pathology protocols.

### Image acquisition and statistical analysis

Slides were imaged via confocal microscopy using a Zeiss LSM-880 (Zeiss, Oberkochen, Germany) system for morphometric analysis, or a Nikon Eclipse/Ds-Ri2 (Nikon, Tokyo, Japan) for quantification analysis. Images were false colored using Fiji. All image analysis and quantification were performed using Fiji (41) in a double-blind fashion and statistical comparisons were analyzed using GraphPad Prism software. Quantification of all images (total “n” and counting metrics) are designated in each figure legend.

## RESULTS

### Morphological changes in the distal colon during development

To understand colon development, we first characterized the morphology of colonic crypts after birth (P1) through adolescence (P21). To accomplish this, we performed Hematoxylin and Eosin (H and E) staining of formalin-fixed, paraffin-embedded tissue sections from wildtype (WT) mice. At P1, the colonic epithelial cells formed a continuous columnar sheet, with occasional folded structures, residing above the non-epithelial cells of the developing lamina propria and muscle tissue (Figure 1A). One week after birth (P7), frequent folds within the epithelium indicated widespread crypt formation (blue arrowheads, Figure 1B). The mucosal and submucosal layers were larger than at P1, the columnar shape of the epithelial cells was more evident, and we detected cells which appear to have similar morphology to adult secretory cells within the epithelial cell layer. Two weeks after birth (P14), the colonic epithelium harbored numerous, deep crypt-like structures with cells organized linearly into a clearly defined crypt-base to crypt-cuff axis. Consistent with the growth from P7 to P17, the mucosa and submucosa were larger (Figure 1C). Lastly, three weeks after birth (P21), the epithelial layer consisted exclusively of crypt-like structures, which contained numerous secretory (yellow asterisks) and absorptive cells (green asterisks). These crypts, as well as the underlying lamina propria and muscle layers, were morphologically indistinguishable from those of adult distal colonic crypts (Figure 1D). In sum, we show the colon undergoes numerous morphological changes as it grows from birth to three weeks of age.

**Figure 1.**
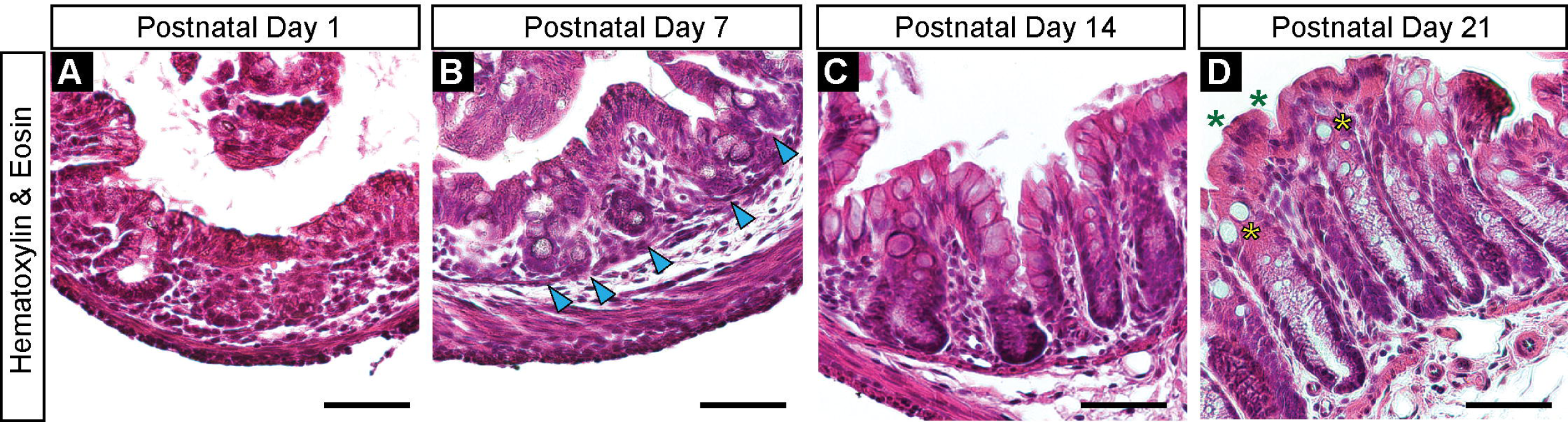
Morphological changes in the distal colon during development. Characterization of the morphology of colonic crypts after birth (P1) through adolescence (P21). Representative colon transverse sections of wildtype mice stained with Hematoxylin and Eosin (A-D). A) At P1, the colonic epithelial cells form a continuous epithelium, populated by columnar cells that reside adjacent to the lamina propria and muscle layers. B) Widespread crypt initiation was detected by the presence of frequent folds within the epithelium at P7 (blue arrowheads). C) At P14, epithelial cell layer was comprised of distinct crypt structures. D) At P21, crypt-like structures are readily apparent, containing secretory and absorptive cells (yellow and green asterisks, respectively). N=5 *C57BL/6* mice surveyed per timepoint. Representative images for each timepoint are shown. Scale bars=50μm.

### Molecular characteristics of the developing colon

In order to better understand the morphological changes we detected from P1-P21, we sought to molecularly characterize the epithelial cells during the same developmental timeframe. To accomplish this, we performed immunostaining of the junctional marker β-catenin, the proliferative marker Kiel-67 (Ki-67), and the stem and progenitor cell marker Leucine-rich repeats and immunoglobulin-like domains 1 (Lrig1). β-catenin is localized to adherens junctions and basal-lateral borders of cells (42). At P1, we observed β-catenin (yellow) localized to epithelial cells lining the mucosal layer and notably, in the small invaginations we had detected by H and E (Figure 2A, A’). At this timepoint, β-catenin was also expressed by non-epithelial cells within the submucosa and muscle tissue (Figure 2A’). At P7, β-catenin was detected in all cells that populate the deep invaginations of developing colonic crypts and was also present within the muscle layer (Figure 2B, B’). The same pattern was detected through P14 and P21 (Figure 2C-D, C’-D’). To visualize proliferation in the developing colon, we assessed protein expression of Ki-67 which is expressed by cells in all active phases of the cell cycle (43). At P1, Ki-67 expression (magenta) was detected throughout the mucosa and submucosa (Figure 2A, A’’) but by P7 and P14, Ki-67 expression was restricted to a subset of cells within both compartments (Figure 2B, B’’, C, C’’). At P21, Ki-67 was largely constrained to the epithelial cells in colonic crypts (Figure 2D, D’’). We next examined Lrig1 expression which is detected on stem and progenitor cells within the adult colon (17, 18). At P1, we observed Lrig1 protein expression (white) restricted to a subset of epithelial cells located within the small invaginations in the mucosa (Figure 2A, A’’’). Lrig1 was also detected in the submucosa, consistent with expression of Lrig1 in the Interstitial Cells of Cajal in the adult colon (44, 45) (Figure 2A, A’’’). As these invaginations enlarge through P7 and P14, Lrig1 expression was confined to cells located near the base of the epithelial folds (Figure 2B, B’’’, C, C’’’). Three weeks after birth, Lrig1 remained restricted to epithelial cells at the base of individual crypts, in a similar expression pattern as is seen in the adult colon (17, 18) (Figure 2D, D’’’). Together, these data describe epithelial cell location, areas of proliferation and expression of Lrig1 in the colon from birth to three weeks of age.

**Figure 2.**
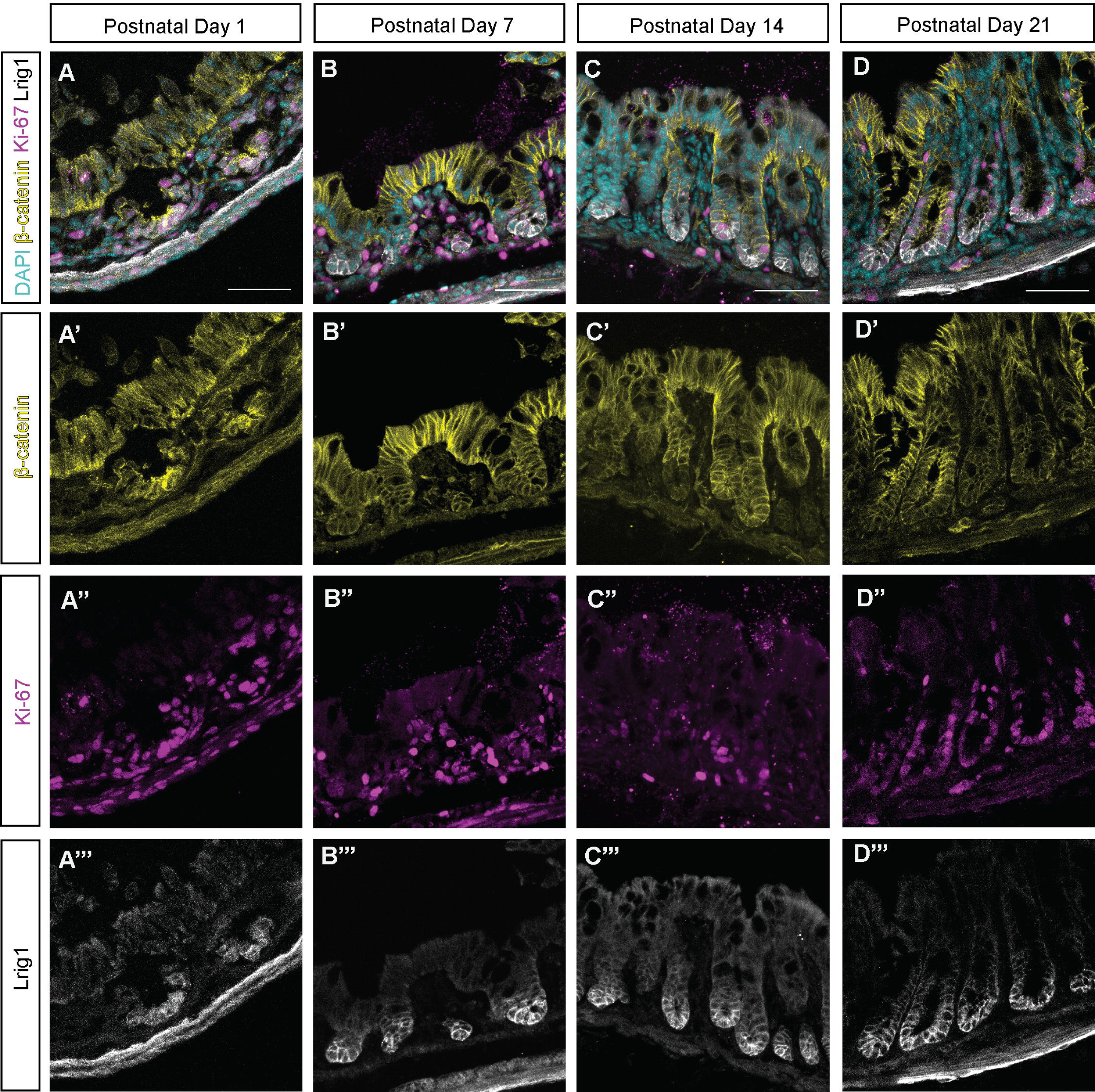
Molecular characteristics of the developing colon. Protein expression of the junctional marker β-catenin, the proliferative marker Ki-67, and the stem and progenitor cell marker Leucine-rich repeats and immunoglobulin-like domains 1 (Lrig1) from P1-P21 (A-D). A’-D’) β-catenin (yellow) detected at cell junctions in the mucosal and submucosal layers. A’’-D’’) Punctate expression of Ki-67 (magenta) detected in the mucosal and submucosal layers. A’’’-D’’’) Lrig1 (white) was detected in cells in both early and late crypt development, as well as in the submucosa. DAPI, (cyan) is marking all nuclei displayed in the merged images (A-D). Images shown are maximum intensity projections of confocal z-stacks taken from 7.5μm to 12μm. N=5 *C57BL/6* mice surveyed per timepoint. Representative images for each timepoint are shown. Scale bars=50μm.

### Lrig1+ cells act as stem cells during colon development

The location of Lrig1 expression throughout colonic epithelial development suggested Lrig1 might be marking adolescent stem cells. We hypothesized if Lrig1 expressing (Lrig1+) cells were acting as stem cells, they may divide and give rise to daughter cells that would populate the crypts. To test this hypothesis, we took a multicolor lineage tracing approach by intercrossing *Lrig1-CreERT2/+* (18) and *R26R-Confetti* (34) mice. The resulting *Lrig1-CreERT2/+;R26R-Confetti* mice received one intraperitoneal injection (IP) of 30mg/kg tamoxifen at P1 to initiate lineage tracing in Lrig1+ cells. We then sacrificed the mice at discrete timepoints to examine fluorescent reporter expression in the colonic epithelium. At P5 we observed multiple populations of Lrig1 lineage-labeled cells (yellow, cyan and red) within the colonic epithelium (Figure 3A). Nearly two weeks later (P17), colon crypts harbored multiple, differently colored cells (polyclonal labeling) or expressed a single color in all cells (monoclonal labeling, Figure 3B). To determine if polyclonal crypts became monoclonal over time, we allowed a cohort of labeled mice to develop to P22 (Figure 3C). At this timepoint, labeled crypts cross-sections were a single color indicating monoclonality. We also detected lineage tracing within the submucosa of the colon at P22 consistent with Lrig1 expression in the Interstitial Cells of Cajal in the adult colon (18, 44). Our lineage tracing data show multiple Lrig1+ cells present at birth can give rise to the epithelial cells in the nascent crypts and these Lrig1+ cells drive clonality within individual crypts over the first three weeks of life.

**Figure 3.**
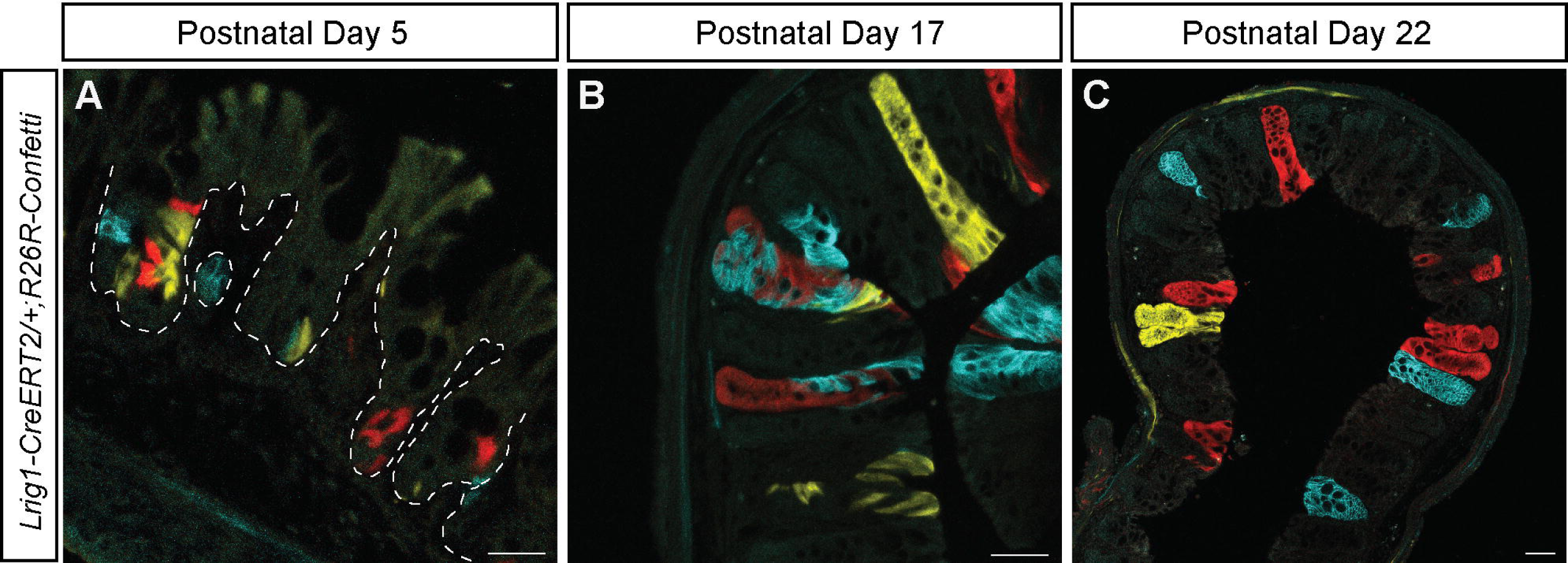
Lrig1+ cells give rise to clonal crypts during colon development. Colon images from *Lrig1-CreERT2/+;R26R-Confetti* mice injected with tamoxifen at P1 and harvested at P5, P17, and P22. A) At P5, multiple populations of Lrig1 lineage-labeled colored cells within the colonic epithelium were detected. White dashed line outlines the epithelial layer. B) At P17, polyclonal (differently colored) crypts and monoclonal (single color) crypts are detected. C) At P22, monoclonal (single color) crypts predominate. N=5 *Lrig1-CreERT2/+;R26R-Confetti* mice per timepoint. Representative images for each timepoint are shown. For A-B, images shown are maximum intensity projections of 12µm confocal z-stacks. For C, 16 confocal images were tiled by the confocal software with 10% overlap in a rectangular grid. Scale bars=50µm.

### Generating the inducible loss of Lrig1 during development

Our data thus far indicate that Lrig1 is expressed in stem cells in the developing colon and we next tested the role of Lrig1 in colon development. In the adult mouse, Lrig1 regulates growth in numerous epithelial tissues (44, 46, 47). During brain development, Lrig1 is also important for regulating cell fate decision making (40). A complete knockout of Lrig1 can result in postnatal death (17), therefore we devised an approach to perform inducible, loss-of-function experiments with the goal of defining the role of Lrig1 without deleting it entirely. To accomplish this, we combined *Lrig1-CreERT2/+* mice (18) with *Lrig1-flox/flox (Lrig1-fl/fl)* mice (EUCOMM; Wellcome Trust Sanger Institute) (48) to create *Lrig1-CreERT2/flox* (*Lrig1-CreERT2/fl)* experimental mice and *Lrig1-WT/flox* (*Lrig1-WT/fl*) littermate controls (Figure 4A). All mice were injected with tamoxifen at P5, which induced recombination of the remaining *Lrig1* allele in the experimental mice (Figure 4B). To examine if this recombination resulted in the loss of Lrig1 protein, we examined *Lrig1-CreERT2/fl* experimental and *Lrig1-WT/fl* control colons for Lrig1 protein expression (17, 18, 46, 49) at specific stages of development (Figure 4C-J). Two days after the induction of recombination (P7), Lrig1 protein (cyan) was uniformly distributed at the base of the epithelial invaginations of the *Lrig1-WT/fl* littermate control colons (Figure 4C), as we observed in WT tissue (Figure 2B, 2B’’’). In our *Lrig1-CreERT2/fl* inducible knockout animals, Lrig1 was virtually undetectable (Figure 4D). The same pattern was observed at P14 for both the control (Figure 4E) and knockout colons (Figure 4F). At P21, Lrig1 was expressed at the base of every control crypt (Figure 4G), as we observed in WT colons (Figure 2D, D’’’). However, in our P21 experimental *Lrig1-Cre/fl* animals, we detected some individual crypts expressing Lrig1 at the crypt-base (yellow asterisk, Figure 4H). In adult colons (P42), Lrig1 was expressed in every control crypt (Figure 4I), as has been reported previously (17, 18), whereas our knockout animals lacked Lrig1 in most crypts, yet we did observe small groups of Lrig1+ crypts (yellow asterisks, Figure 4J), similar to a mosaic pattern. Given this expression of Lrig1 in discrete cells within our experimental colons, we then revisited the same experimental images from Figure 4D, F, H and J to test if enhancing the brightness of Lrig1 in the image would allow us to detect any Lrig1+ cells at P7 and P14. With these enhanced images, we detected single Lrig1+ cells (white arrowheads, Figure 4 D’ and F’) at the P7 and P14 timepoints. The same groups of Lrig1+ cells present in the P21 and P42 experimental colons were also more readily visible with increased brightness (yellow asterisks, Figure 4H’ and J’). In sum, our inducible knockout approach generated efficient knockout of Lrig1 as early as two days post-induction (P7) and this was maintained through P14. At the later timepoints, Lrig1 expression was readily detectable in the *Lrig1-CreERT2/fl* inducible knockout mice, therefore we focused on the P7 and P14 timepoints to evaluate the impact of the loss of Lrig1.

**Figure 4.**
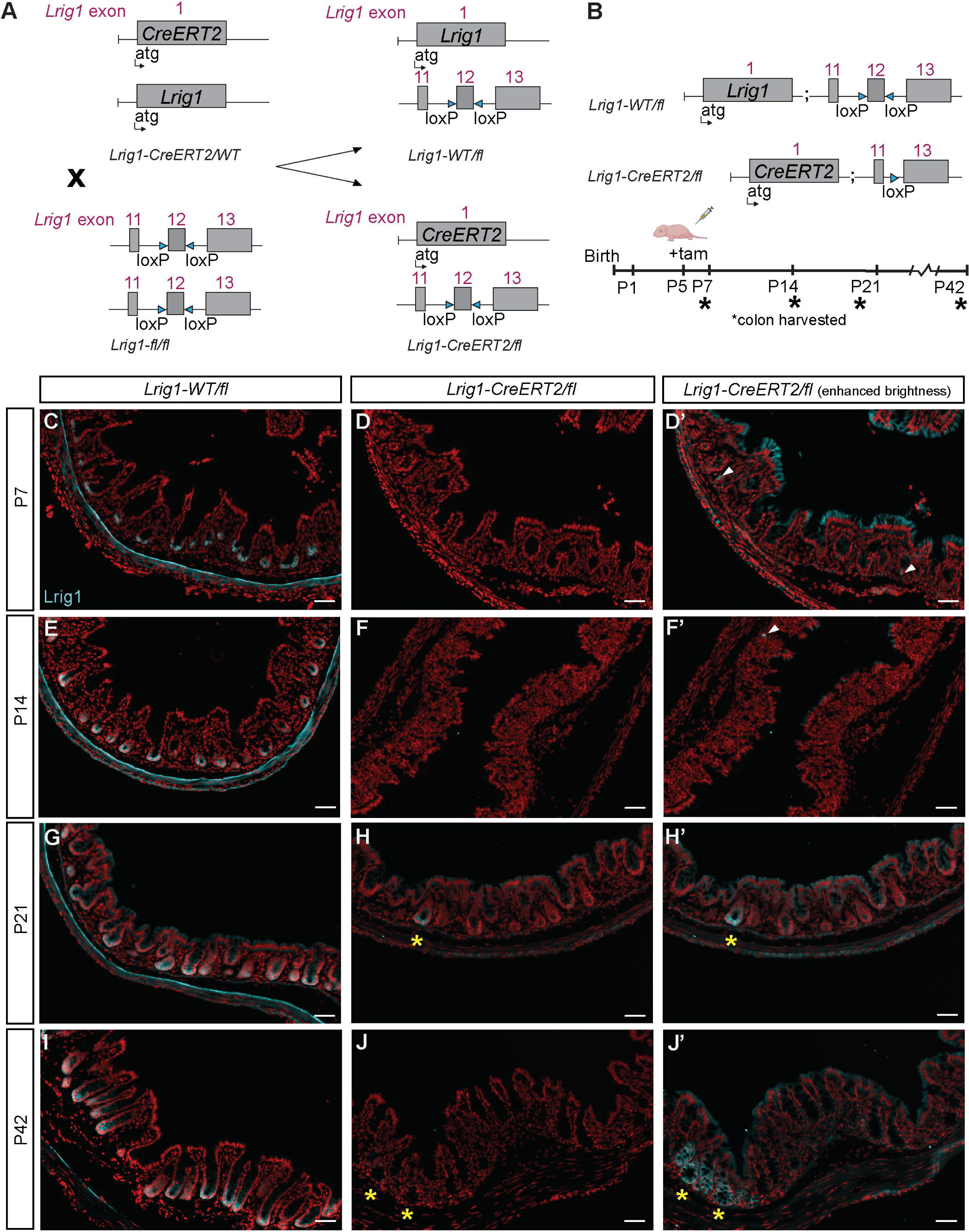
Generating the inducible loss of Lrig1 during development. Colon images from *Lrig1-CreERT2/fl* mice injected with tamoxifen at P5 and harvested at P7, P14, P21, and P42. A) Schematic for the generation of *Lrig1-CreERT2/fl* experimental and *Lrig1-WT/fl* littermate control mice. B) Experimental schematic for the injection of *Lrig1-CreERT2/fl* experimental and *Lrig1-WT/fl* littermate control mice at P5, and colon tissue harvested at P7, P14, P21, and P42 (asterisks). C-J’) Epifluorescent images displaying protein expression of Lrig1+ cells (cyan) and DNA (DAPI; red) at the timepoints surveyed. D’, F’, H’ and J’) Enhanced images by increasing the brightness of D, F, H and J images, to showcase Lrig1+ cells (white arrowheads) and Lrig1+ crypts (yellow asterisks) in *Lrig1-CreERT2/flox* mice. N=6 mice per genotype per timepoint. Representative images are shown. Scale bars=50μm.

### Lrig1 restrains proliferation in developing colonic crypts

Lrig1 is a known regulator of growth and proliferation(18, 44, 47, 50, 51). In addition, *Lrig1* complete knockouts have significant exogenous proliferation within the intestine before their postnatal death (17) and expression of Lrig1 is rapidly expanded in the colon as a part of the regenerative process after injury to the epithelium (21). Given these data, we hypothesized Lrig1 regulates proliferation during colon development and loss of *Lrig1* in our inducible knockout mice would result in hyperproliferation. In Figure 1, we show widespread crypt formation and cell proliferation occurred during the first week of birth (Figure 1B), so we next examined whether loss of *Lrig1* affected cell proliferation in the *Lrig1-CreERT2/fl* inducible knockout mice compared to our *Lrig1-WT/fl* littermate control animals at P7 (Figure S1). To assess proliferation, we examined the P7 colonic epithelium for the expression of the proliferative marker Ki-67. After analyzing both cohorts of animals, we detected indistinguishable patterns of Ki-67 expression between our injected inducible knockout mice, two days after induction, compared to their littermate controls (Figure S1A-B’). We further quantified this observation by counting the total Ki-67+ cells per 100µm^2^ of total mucosa area (Figure S1C). As colon proliferation is ongoing but moderate from P8 to P16, compared to earlier developmental timepoints (14), we reasoned that examining proliferation at P14 in our knockout *Lrig1-CreERT2/fl* and *Lrig1-WT/fl* littermate control animals, would be an appropriate timepoint to measure any potential increase in proliferation in the loss-of-function animals. In agreement with ^3^[H] assays (14), Ki-67 protein was detected in a few, discrete cells at the base of colonic crypts in our littermate controls (*Lrig1-WT/fl* mice; Figure 5A-A’). This was consistent with what we observed in WT crypts at this same developmental stage (Figure 2C, C’’). In contrast, we detected expanded Ki-67 expression within the crypts of our *Lrig1-CreERT2/fl* mice (Figure 5B-B’). We quantified this observation across all mice, as in Figure S1, and show proliferation was significantly increased in our P14 knockout mice, compared to the littermate controls (Figure 5C, *p*<.0001, n=6 for each genotype). As experimental *Lrig1-CreERT2/fl* mice displayed a mosaic pattern of Lrig1 expression at P21 (shown in Figure 4H’), we hypothesized crypts retaining Lrig1 would have less proliferation than knockout crypts from the same animal. To test this, we examined Ki-67 protein expression in Lrig1-expressing and Lrig1-missing adjacent crypts (Figure 5D-D’’’). In agreement with our hypothesis, we detected a significant decrease in the number of Ki-67+ cells in Lrig1-expressing crypts compared to their Lrig1-missing adjacent counterparts, in many of the mice we examined (Figure 5E).

**Figure 5.**
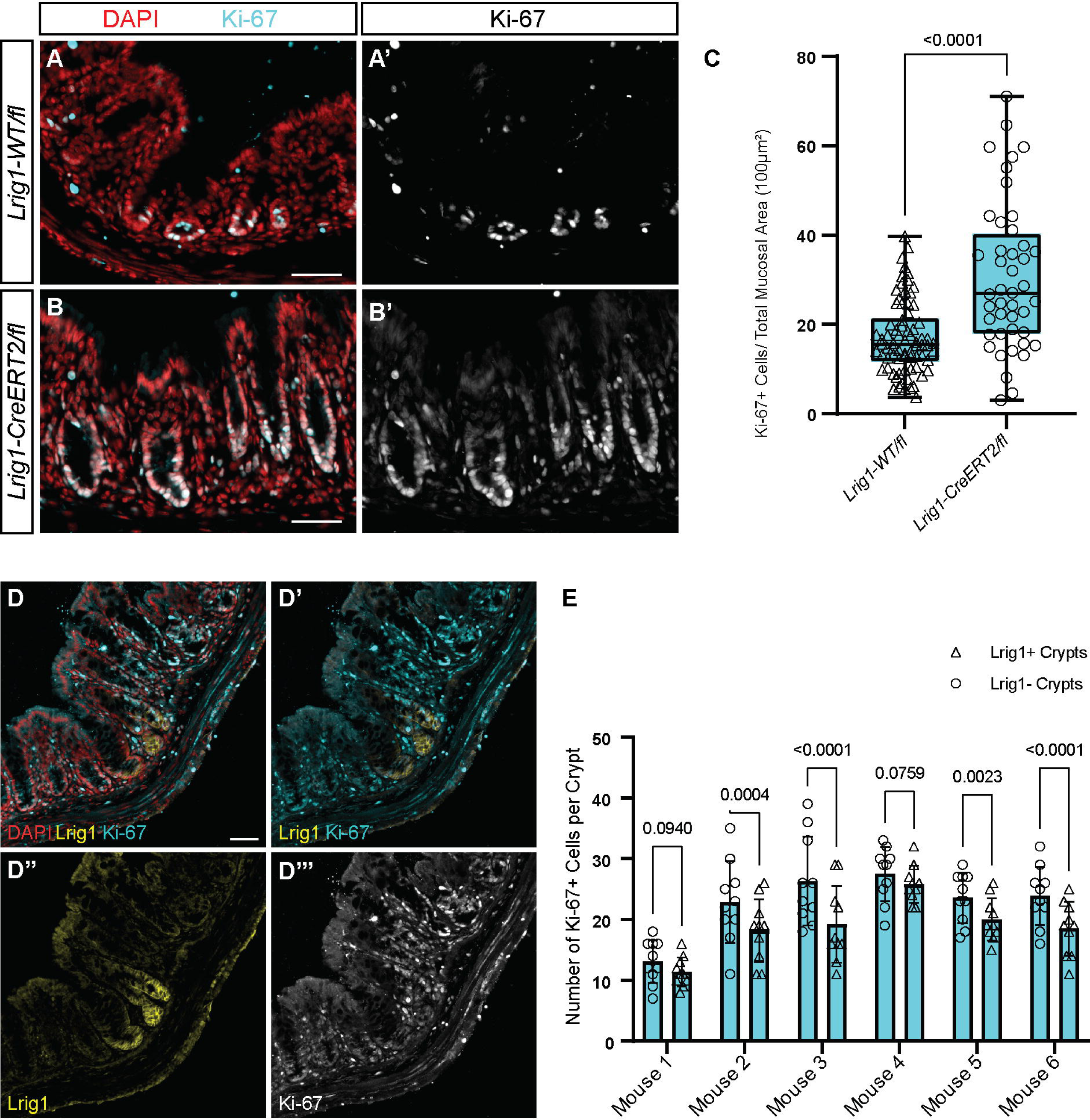
Lrig1 restrains proliferation in developing colonic crypts. Colon images from *Lrig1-CreERT2/fl* experimental and *Lrig1-WT/fl* littermate control mice injected with tamoxifen at P5 to induce the loss of Lrig1, and tissue was harvested at P14 (A-C) and P21 (D-E). A-B’) Proliferation detected by expression of Ki-67 (cyan) expression in *Lrig1-CreERT2/fl* experimental and *Lrig1-WT/fl* littermate control mice. All cells present are indicated by the nuclear marker DAPI (red). C) Quantification of Ki-67+ cells per total mucosal area (100μm^2^). N=6 mice per genotype, 110,000µm (+/-15k µm) mucosal area analyzed per mouse. Box-whisker plot represents individual data points with whiskers representing minimum to maximum values. Significance was determined by an unpaired *t*-test and *p* values are indicated. D-E) Analysis of Lrig1 and Ki-67 expression in mosaic crypts D-D’’’) Protein expression of Lrig1 and Ki-67analyzed in *Lrig1-CreERT2/fl* experimental mice displaying representative adjacent Lrig1-expressing and Lrig1-missing crypts. D) Composite image of DAPI (red), Lrig1 (yellow), and Ki-67 (cyan). D’) Overlay of Lrig1 and Ki-67 protein expression. D’’) Single channel of Lrig1 protein expression. D’’’) Single channel of Ki-67 protein expression. E) Quantification of number of Ki-67+cells per crypt in n=6 *Lrig1-CreERT2/fl* mice. Bar graph represents individual data points in Lrig1-expressing and Lrig1-missing crypts with error bars representing the standard error of the mean. Significance was determined by a nested paired *t*-test and *p* values are indicated. For A-E, images are epifluorescence images. Scale bars=50μm.

### Loss of Lrig1 does not impact differentiation

In the brain and the skin, Lrig1 controls a balance between proliferation and differentiation (38–40). As the loss of Lrig1 increased proliferation in the developing colon at P14, we next wanted to examine if the loss of Lrig1 inhibits differentiation of colon epithelial cells at the same developmental timepoint. To examine this, we performed immunofluorescent antibody staining for common differentiation markers of both secretory and absorptive cell types in our *Lrig1-CreERT2/fl* inducible knockout mice and *Lrig1-WT/fl* littermate controls (Figure 6). We used Mucin2 (Muc2, cyan) (52) to detect emerging secretory cells at P14 (Figure 6A-B) and compared the two genotypes. We detected no quantitative difference in Muc2 expression between the genotypes both qualitatively and quantitatively (Figure 6C). To detect absorptive cells, we used the brush border marker phalloidin (cyan) (53) and we detected no qualitative difference in phalloidin expression between the genotypes (Figure 6D-E’). We next examined the presence of emerging enteroendocrine and Tuft cells by staining with Chromogranin A (CgA, cyan, Figure 6F-G’) (26, 54) and Doublecortin-like kinase 1 (Dclk1, cyan, Figure 6H-I’) (55), respectively. As with Muc2 and phalloidin, there was no obvious difference in the differentiation pattern of these cells in both genotypes (Figure 6 F-I’). In sum, these data show loss of *Lrig1* does not impact colon epithelial cell differentiation during the developmental time window from P5 to P14.

**Figure 6.**
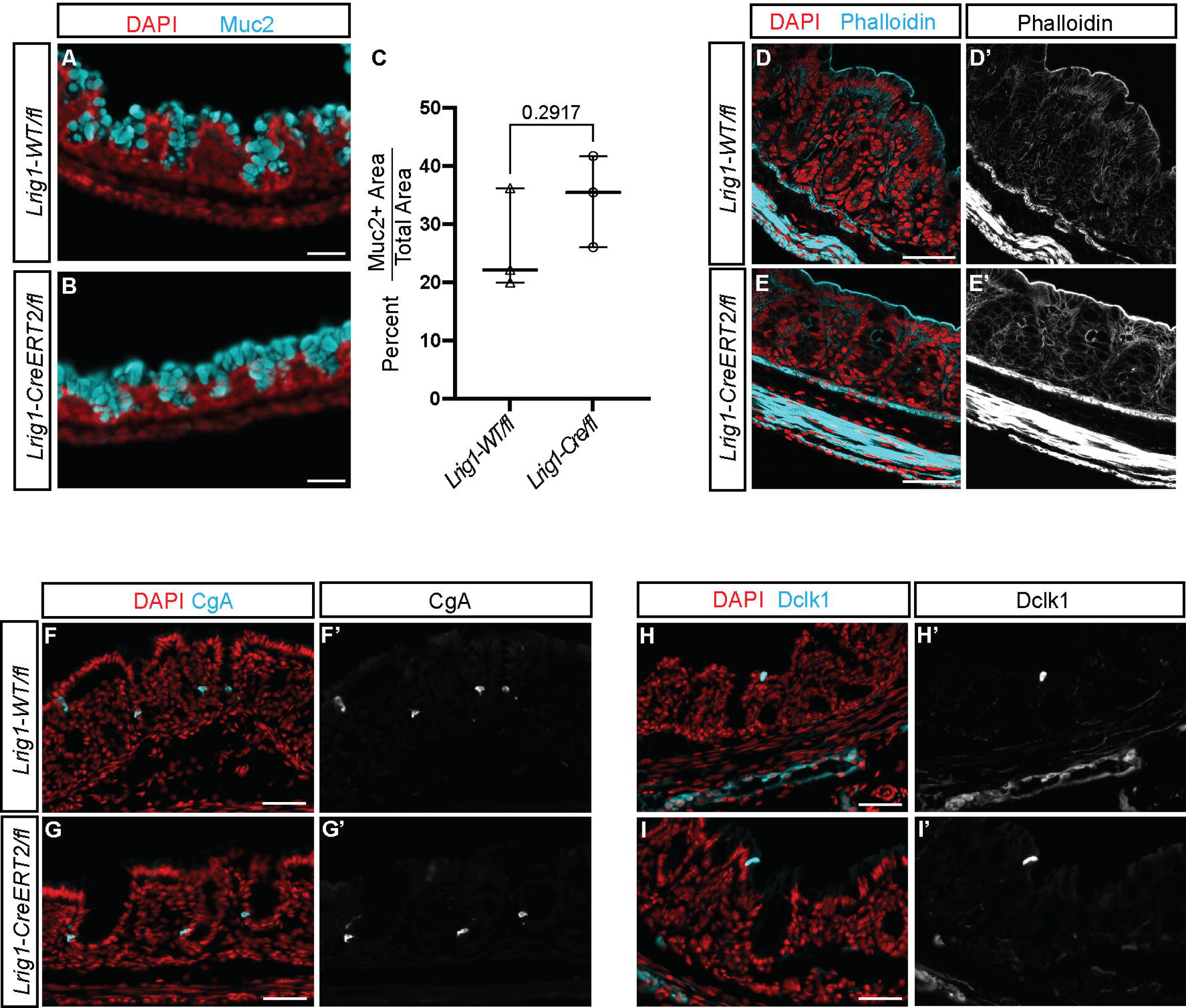
Lrig1 does not regulate differentiation in developing colonic crypts. Protein expression of the goblet cell marker Mucin 2 (Muc2), enteroendocrine cell marker Chromogranin A (CgA), tuft cell marker Doublecortin-like kinase 1 (Dclk1), and absorptive cell marker phalloidin in *Lrig1-CreERT2/fl* experimental and *Lrig1-WT/fl* littermate control mice. For all images, the protein of interest is labeled in cyan, and all nuclei are labeled with DAPI in red. A-B) Epifluorescence images of Muc2 protein expression in *Lrig1-CreERT2/fl* and *Lrig1-WT/fl* mice. C) Graph displaying percent positive Muc2 area per total epithelial area for *Lrig1-CreERT2/fl* and *Lrig1-WT/fl* mice. For the whisker plot, the individual data points represent the mean percent positive Muc2 area per total epithelial area for each mouse. N=3 mice and 10 random images were acquired per mouse for each genotype. Significance was determined by an unpaired *t*-test and the *p* value is indicated. D-E’) Expression of F-actin detected by the presence of phalloidin (cyan) using confocal microscopy (single slice shown). F-G’) Epifluorescence images of Chromogranin A (CgA; cyan) protein expression in *Lrig1-CreERT2/fl* and *Lrig1-WT/fl* mice. H-I’) Epifluorescence images of Doublecortin-like kinase 1 expression (Dclk1; cyan) protein expression in *Lrig1-CreERT2/fl* and *Lrig1-WT/fl* mice. All images are representative and acquired from P14 colon. Scale bars=50μm.

## DISCUSSION

Our research is one of the first to define the morphological and molecular characteristics of the developing distal mouse colon three weeks after birth. Our data describes three key molecular characteristics of the developing colon: epithelial cell location, areas of proliferation, and emergence and expression of a stem and progenitor cell marker, Lrig1. Using a novel *Lrig1-CreERT2/+;R26R-Confetti* reporter mouse, we demonstrate that Lrig1+ cells are present at birth and these cells can give rise to colonic crypts. To test the role of Lrig1 in development, we created an Lrig1 inducible knockout mouse (*Lrig1-CreERT2/fl)* and after knocking out Lrig1 at discrete timepoints after birth, our data indicate that inducible loss of Lrig1 leads to hyperproliferation two weeks after birth. In addition, we show this phenotype does not impact differentiation in the developing colon. In sum, our data is the first to show Lrig1 is required to restrain proliferation during colon crypt development, specifically during the first two weeks after birth and unlike the adult colon, hyperproliferation does not lead to a loss of differentiation during colon development.

The first aim of our study was to morphologically characterize the development of distal colonic crypts using modern pathology staining techniques. In addition, despite the numerous distinctions between the colon and the small intestine, most of the research conducted on the developing intestine has historically been focused on the small intestine (16, 22, 56–59) and does very little to address colon development. Therefore, we thought it was pertinent to examine the formation of colonic crypts as colon development proceeds. Our data demonstrate distal colonic crypt formation is widespread by the first postnatal week and hallmarks of epithelial cell differentiation can be observed by the second postnatal week. During this early timeframe, the distal colon is already comprised of cells displaying morphological features of classically differentiated cells While the purview of our study was kept to a relatively small developmental time window, several interesting areas for further research arise from these morphological data. In Figure 1 we show small invaginations, which represent nascent crypts, are present the day after birth. Going forward, it will be important to examine the formation of the colon from the late embryonic days up until birth to map tube development and observe the cellular organization that occurs to generate these nascent crypts. It will also be interesting to examine whether or not this crypt formation in the colon mimics the same process as the small intestine. Our data from P21 mice indicate that the adolescent distal colonic crypts present appear indistinguishable from adult crypts, however testing whether they are truly adult-like in nature, in terms of size, width, and function will be crucial to our understanding of colonic function during development. Finally, a careful examination of crypt bifurcation and expansion of the crypts during this time frame will promote our understanding of how the colon crypts distribute themselves along the rostral to caudal axis.

The extensive morphological changes we show in Figure 1 are accompanied by the molecular marker expression shown in Figure 2. Our molecular characterization illustrates the tight epithelial organization, the highly proliferative epithelial cells present, and the expression of the stem and progenitor marker Lrig1, throughout colon development. Most of the previous studies examining the cellular characteristics of the developing colon were performed before advanced molecular markers were developed and a clear next step is to more fully examine additional molecular features of developing colonic crypts. These analyses could include several different avenues of investigation; one path that might be particularly interesting would be examining the emergence of stem cell support populations during crypt emergence and expansion. Studies like these will help us understand how these cells come to reside next to each other in adult homeostasis and if they have a symbiotic relationship, as in adult colonic tissue. In the developing small intestine, stem cell support cells called Paneth cells emerge during the first week after birth and a dramatic increase in Paneth cell number occurs between two and four postnatal weeks (60). Paneth cells produce factors that promote stem cell homeostasis in the small intestine (61) and are important for the process of crypt expansion (62). As Paneth cells do not exist in the mouse colon, cells marked with Regenerating Family Member 4 (Reg4) have been defined as deep crypt secretory cells, and often act as Paneth cell equivalents (60, 63). It is currently unknown when Reg4+ cells emerge during development and if they are required for developmental stem cell homeostasis, or crypt growth. This will be an important area for future investigation and could be accomplished by combining any number of colon stem cell fluorescent reporters (25, 45, 64, 65) with the *Reg4-dsRed-DTR* mouse to examine patterning and crypt formation in the presence or absence of Reg4+ cells. Certainly, the possibilities for expanding our molecular knowledge in colon development are numerous and could also include examining of the emergence of cellular transporters, junctional proteins, and additional differentiation markers. Ultimately, discovering the molecular underpinnings of how colonic crypts develop may be informative for our understanding of colonic disease progression and crypt regeneration in humans.

Our study is the first to use multicolor lineage tracing in Lrig1+ cells throughout colon development to examine the clonality potential of young epithelial stem cells. Lrig1 expressing (Lrig1+) cells are present at P1 in the base of immature crypt-like folds and using our multicolor approach, we demonstrate these Lrig1+ cells continuously produce daughter cells, which give rise to fully labeled, clonal crypts by P22. It is intriguing that multiple populations of Lrig1+ cells establish clonal crypts three weeks after birth, as this is a distinct timeline from clonality studies in adult mice (34). We show Lrig1+ cells can be detected as early as P1 and are localized to the base of nascent and developing crypts throughout colon crypt development, suggesting that Lrig1+ cells may be involved in crypt establishment. In the future it will be important to define where the Lrig1+cells arise from, both in cellular lineage and in anatomical location. Answering these questions may give us a fuller picture of the origins of these cells, which are critical for appropriate proliferation in developing crypts.

Our inducible, loss-of-function approach for eliminating Lrig1 enabled us to circumvent the birth defects associated with the Lrig1 knockout (17) and ultimately allowed us to define the molecular function of Lrig1 during colon development. Loss of Lrig1 resulted in increased proliferation in developing colonic crypts two weeks after birth (P14), yet our data also revealed populations of Lrig1+ “escaper cells” detected at P7 and P14 and Lrig1+ “escaper crypts” were detected and persist one week later (P21). These data illustrate a selection against Lrig1 elimination during colon crypt development and a dominant push for Lrig1 expression as crypts are forming. Examining the molecular regulators of this selection towards homeostasis will be important. Likely, this push is dictated by a multitude of signaling mechanisms which regulate a balance between proliferation and differentiation, as in adult mice (61, 61, 63, 66–70). To this end, we show Lrig1 suppressed proliferation in a narrow developmental window, yet this level of proliferation can vary, even between adjacent crypts. It is currently unknown how Lrig1 is regulating proliferation during colon development, but studies from the small intestine may offer clues. It is well-established that the Notch and Wnt cell signaling cascades regulate epithelial cellular proliferation and differentiation (71–74) in the adult small intestine and colon. In addition, an interplay between Wnt, BMP, and Hedgehog signaling cascades are important for crypt formation in the small intestine (57, 75, 76). While we know Lrig1 is important for regulating proliferation, we do not yet know where Lrig1 fits in between these signaling cascades. Further studies examining the cellular mechanisms impacted due to both the loss and overexpression of Lrig1, will be informative to help us build a molecular model for colon crypt development.

Perhaps the most striking observation from our developmental studies is the hyperproliferation we observe in our *Lrig1-Cre/fl* experimental mice at P14 does not result in a loss of differentiation, as commonly seen in adult mice (77, 78). Our data indicate that both absorptive and secretory cell populations are still present in the highly proliferative epithelium, suggesting these differentiated cells are actively proliferating. As these differentiated cells do not proliferate in adult tissues, our data support a model where molecular signals may govern newly differentiated cells in the developing colon differently, compared to those which regulate the rapidly renewing adult intestinal epithelium.

In sum, our studies clearly define the importance of studying Lrig1 and its role in colon development. We address a critical gap in the intestinal development literature and provide new information about the molecular cues that guide colon development. Using a novel inducible knockout of Lrig1, we show Lrig1 is required for appropriate colon epithelial growth and illustrate the importance of Lrig1 in the establishment of developing colonic crypts.

## ACKNOWLEDGEMENTS

We thank our colleagues at the University of Oregon for the critical reading of this manuscript. We also thank the Kryn Stankunas lab for sharing reagents and the donation of the Confetti mice for our study and Astra Henner for her technical expertise. We thank Adam Fries from University of Oregon’s Genomics & Cell Characterization Core imaging facilities for his imaging expertise.

## AUTHOR CONTRIBUTIONS

RH designed, executed, and analyzed the study and wrote the manuscript. NJ provided technical assistance, mouse expertise and manuscript feedback. AZ designed, executed, and analyzed the study and wrote the manuscript.

---

**Figure S1.**
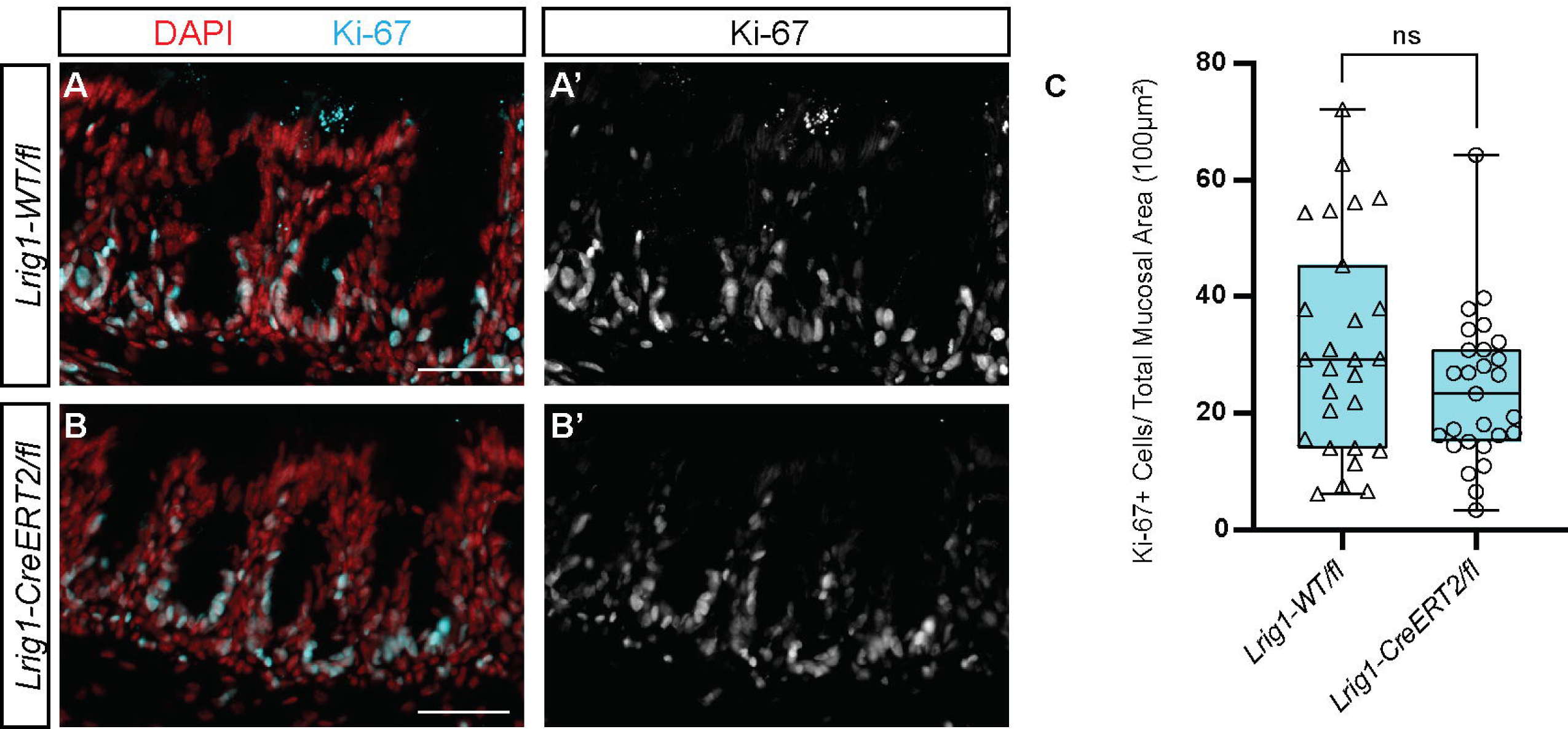
Loss of Lrig1 does not affect proliferation at P7. Colon images from *Lrig1- CreERT2/fl* experimental and *Lrig1-WT/fl* littermate control mice injected with tamoxifen at P5 to induce the loss of Lrig1, and tissue was harvested at P17 (A-C). A-B’) Proliferation detected by expression of Ki-67 (cyan) expression in *Lrig1-CreERT2/fl* experimental and *Lrig1-WT/fl* littermate control mice. All cells present are indicated by the nuclear marker DAPI (red). C) Quantification of Ki-67+ cells per total mucosal area (100μm^2^). n=6 mice per genotype, 110,000µm (+/-15k µm) mucosal area analyzed per mouse. Box-whisker plot represents individual data points with whiskers representing minimum to maximum values. Significance was determined by an unpaired *t*-test and *p* values are indicated. All images are epifluorescence images. Scale bars=50μm.

